# Mechanism of upstream promoter element stimulation of transcription at a ribosomal RNA promoter determined by single-molecule imaging

**DOI:** 10.1101/2020.02.17.953182

**Authors:** Jeffrey P. Mumm, Larry J. Friedman, Jeff Gelles

## Abstract

DNA elements upstream of transcription promoters play a role in regulating transcription initiation in all organisms. In bacteria, upstream A-T rich sequences called UP elements can stimulate transcription through contact with the α subunit C-terminal domain (αCTD) of core RNA polymerase (RNAP), but the kinetic mechanisms by which they do so remain unclear. We investigated the role of the UP element in stimulating initiation from the strong E. coli 16s rRNA promoter using single-molecule fluorescence microscopy to visualize σ^70^RNAP holoenzyme binding and the formation nascent RNA by oligonucleotide probe hybridization on individual DNA molecules containing the *rrnB* P1 promoter. By directly detecting initial binding of σ^70^RNAP to promoter and monitoring the lifetimes of promoter-polymerase complexes, the experiments reveal the kinetic mechanism of polymerase recruitment to the promoter and the subsequent conformational change that stabilizes binding. The presence of UP stimulated the rate of initial binding of polymerase to promoter by at least six-fold, and this stimulation was fully sufficient to account for the increase in initiation rate by UP. Thus, UP likely functions at this strong promoter simply by acting as a binding target for the rapidly reorienting αCTD domain tethered to the core polymerase. In contrast, there were only minor effects of UP on the measured rates of the conformational change or the dissociation rates of the initial σ^70^RNAP promoter complexes. These studies define a paradigmatic kinetic mechanism for stimulation of transcription initiation by direct αCTD-DNA interactions. This mechanism can serve as a building block of more complex regulatory architectures in which αCTD promotes transcription through interactions with both DNA and protein activators.

## Introduction

Multi-subunit RNAPs initiate the transcription of DNA through a process that is central to the regulation of gene expression in all domains of life [1–3]. A quantitative understanding of this process is vital in outlining the bacterial regulatory network and its dynamic response to environmental changes, identifying and characterizing new targets for antibacterial drug design [4], and developing baseline kinetic mechanisms of transcription initiation to maximize the information gained from genome-wide comparison experiments [5].

The *E. coli rrnB* P1 promoter drives transcription of the 16S rRNA and as such is one of the most actively transcribed bacterial promoters driven by the σ^70^RNAP holoenzyme during exponential growth of cells [6]. In starvation, however, reduced concentration of the initiating NTP and an increased concentration of the alarmone nucleotide ppGpp induce a large decrease in initiation at this promoter. Despite its high activity, *rrnB* lacks many of the hallmarks of strong σ^70^-dependent promoters: it has a non-consensus −35 hexamer, a non-extended −410 box, and a non-ideal −10/−35 spacer. Mutation of these non-ideal elements to an ideal sequence alleviates growth rate-dependent control at the *rrnB* P1 locus [7, 8], consistent with the hypothesis that instability of σ^70^RNAP complexes with the core promoter plays an essential role in regulation.

In addition to the core promoter elements, *rrnB* P1 also contains an upstream promoter (“UP”) element at positions −65 to −41 relative to the transcription start site. Mutations in UP dramatically reduce initiation frequency both *in vivo* [9–12] and *in vitro* [11]. Interestingly, UP ablations can also diminish growth-rate dependent regulation of rRNA promoters [7, 8]. The presence of sequence elements upstream of a promoter that affect promoter activity and regulation is a common theme in both prokaryotes and eukaryotes. In general, such sequences may serve as targets for transcription factor binding or may interact directly with the core transcriptional machinery or both. At some promoters, UP elements may be the promoter regions that make earliest contact with the polymerase during initiation [1, 13, 14]. The UP element is thought to act at least in part through direct binding of the flexibly tethered carboxy-terminal domains of the α subunits of RNAP (α-CTD) [11, 15, 16] in a manner similar to a class I transcriptional activator [16, 17]. This interaction is thought to increase the populations of some types of RNAP-promoter complexes [9, 18], but the kinetic mechanisms by which this is achieved have been challenging to decipher, in part because UP may act at multiple steps in initiation [9, 19–23].

Here we investigate the roles of the UP element on the kinetic processes required for initiation at the *rrnB* P1 promoter. The experiments utilized the colocalization single molecule spectroscopy (CoSMoS) technique previously used to define the kinetic mechanism for initiation at the *E. coli* σ^54^ promoter *glnA* [24, 25] and the mechanism of rrnB repression by secondary channel factors GreB and DksA [26]. This and other single-molecule techniques used to study transcription initiation (see refs. [27–29] and refs. therein) have a number of advantages over traditional ensemble biochemical techniques that include the abilities to discriminate between intermediates from an asynchronous mixture of complexes, to characterize forward and reverse reaction steps in a single experiment, and to directly observe initial RNAP association with a promoter DNA.

## Results

### Observing σ^70^RNAP-*rrnB* P1 Complex Formation and Transcription Initiation on single DNA molecules

To examine the role that UP plays in initiation at *rrnB* P1, we constructed a transcription template containing the wild type *E. coli rrnB* P1 promoter and 251 bp of downstream sequence (WT; Fig. 1A). We also made control templates UP^−^, P1^−^, and UP^−^P1^−^ in which the UP element, the −35 and −10 box core promoter elements, or both were mutated. All four DNA variants also contain the less-active *rrnB* P2 promoter [30] 117 nt downstream of the P1 position.

**Figure 1.**
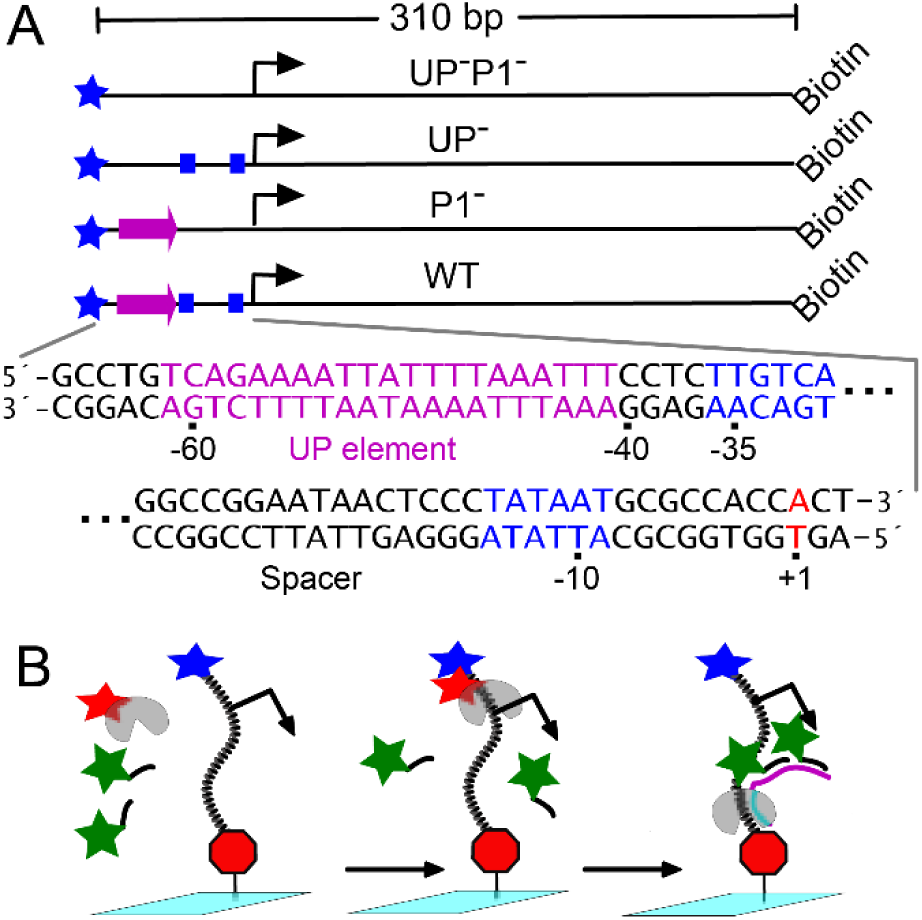
Design of the transcription templates and the single-molecule transcription experiments. **(A)** Templates and control DNAs. The WT template consists of the promoter and initial transcribed region of *E. coli rrnB* P1. The promoter includes UP element (magenta), core promoter elements (−35 and −10 boxes; blue), and transcription start site (bent arrow and red nucleotide). Other DNAs were identical except that one or more promoter elements were ablated by mutation. DNAs were 5’- end labeled with AlexaFluor 488 (blue star) and biotin. **(B)** Experiment design. DNA was tethered to the flow chamber surface (cyan) by avidin (red octagon). The solution introduced into the chamber contained NTPs, *E. coli* σ^70^-RNAP (gray) incorporating Cy5 (red star)-labeled σ^70^, and two Cy3 (green star) hybridization probe oligonucleotides complementary to sequences near the 5′-end of the nascent transcript (magenta). σ^70^RNAP binding was detected as appearance of a red σ^70^-RNAP fluorescent spot at a position where a blue template spot was observed; transcript production is detected as appearance of a green probe spot at the blue template spot position.

We used the CoSMoS technique [24, 26, 31–34] to directly observe binding of single RNAP holoenzyme (σ^70^RNAP) molecules to the WT DNA and the production of individual transcript molecules by these promoter-holoenzyme complexes. WT DNA molecules were tethered at their downstream ends to the surface of a coverslip flow chamber via a biotin-avidin linkage (Fig. 1). The upstream end was labeled with a blue-excited fluorescent dye. The chamber was monitored using total internal reflection fluorescence (TIRF) microscopy, in which only dye moieties that are tethered to the surface are visualized as discrete spots. The surface density of the DNA was kept sufficiently low that the images showed well defined, isolated spots of fluorescence (100-200 per field of view) corresponding to individual DNA molecules (Fig. 2A, left).

**Figure 2.**
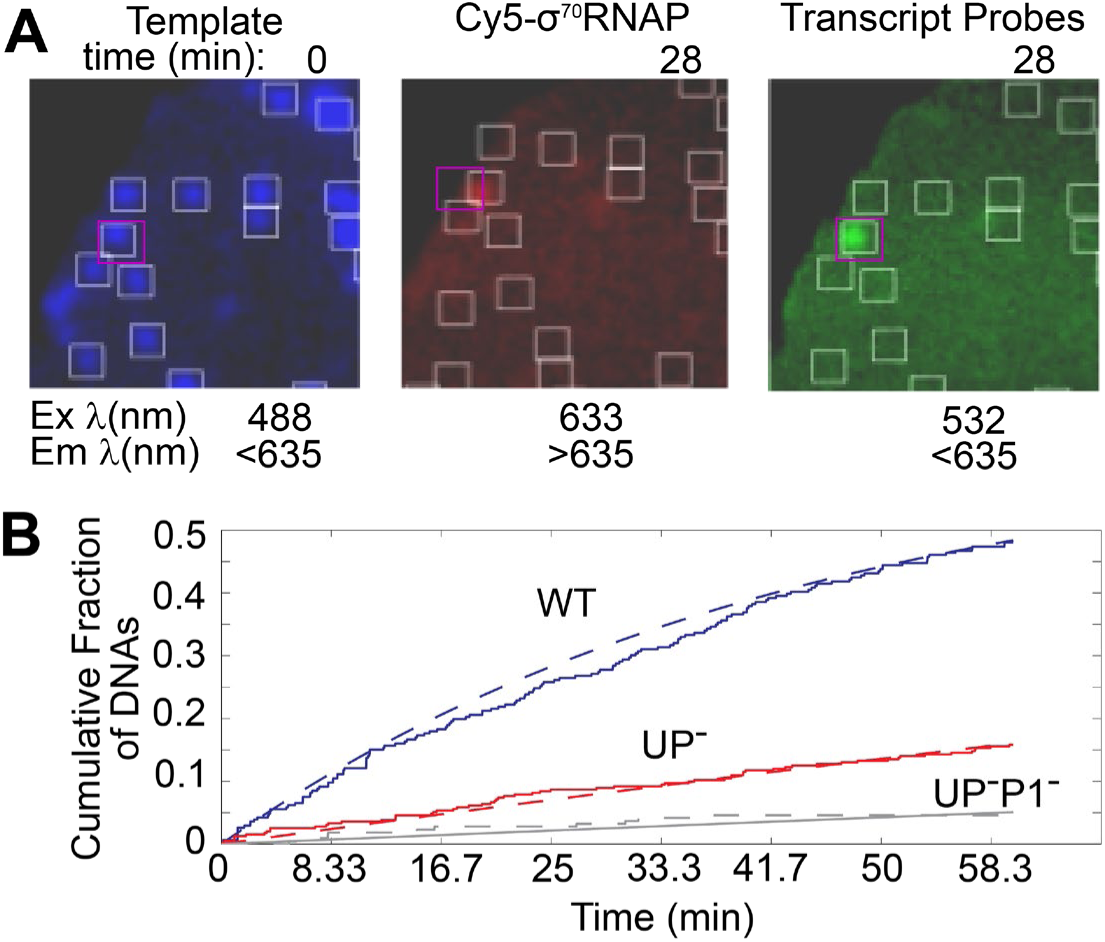
Effect of UP on transcription initiation on single *rrnB* P1 DNA molecules. **(A)** Pseudocolor images (5.4 × 5.4 µm) of a portion of the microscope field of view imaged at excitation (Ex) and emission (Em) wavelengths corresponding to the template, Cy5-σ^70^RNAP, and transcript probe. Superimposed squares mark the positions of spots in the left image and the corresponding positions in the center and right images. Images were recorded at the indicated time after addition of 0.75 nM Cy5-σ^70^RNAP and NTPs. **(B)** Cumulative fraction of DNA molecules on which transcript probe had appeared by the indicated time (solid lines). Exponential fits (dashed lines; see Methods) yielded transcription initiation rates of (3.8 ± 0.8) × 10^−4^ s^−1^ (S.E.) (WT DNA; blue; *N* = 443), (0.49 ± 0.1) × 10^−4^ s^−1^ (UP^−^ DNA; red; *N* = 392), and (0.15 ± 0.002) × 10^−4^ s^−1^ (UP^−^P1^−^; black; *N* = 216). Imaging conditions: 8 s per frame; 25 µW 532 nm excitation.

Next, we introduced a solution containing σ^70^RNAP, labeled with a red-excited Cy5 dye on the σ^70^ subunit [35], and the four nucleoside triphosphates (NTPs). The solution also contained two green-excited dye labeled oligonucleotide probes complementary to the transcript sequence at positions 1-26 and 27-54 (Table S1). Spots of Cy5-σ^70^RNAP fluorescence appeared on the surface (Fig. 2A, center). Typically, 96% of these σ^70^RNAP spots colocalized with recorded positions of individual DNA molecules, suggesting that the spots represent polymerase bound to DNA. Some σ^70^RNAP spots persisted for the full duration of the recording (∼1 hour); others eventually disappeared. Spots that were observed to disappear typically did so completely in a single frame interval, indicating that single σ^70^ subunit was present and then either dissociated from the complex or had its single dye label photobleached by the excitation laser.

Over the course of the recording, spots of green fluorescence (transcript probe) were also observed to colocalize with DNA positions, indicating hybridization with an RNA produced by transcription of the template (Fig. 2A, right). Probe fluorescence intensity was sometime observed to appear in two discrete steps, suggesting that we can detect hybridization to both target sequences on the transcript. Even though elongation of the 240 nt transcript is expected to take <30 s under these conditions, many of these probe spots persisted for tens of minutes, consistent with the expectation that avidin linked to the downstream end of the template will act as a roadblock to movement of the transcript elongation complex (TEC) and cause it to stall [36, 37].

To check that colocalization of transcript probe spots with DNA molecules were the result of σ^70^RNAP-mediated transcription events, we compared the results from the experiment with all four NTPs to a control performed in the absence of GTP. Over a 3,600 s course of observation, 48% (148 of 306) DNA locations exhibited colocalized probe spots in the former, compared to only 1% (2 of 161) in the control. Moreover, in transcription experiments containing all four NTPs with the UP^−^P1^−^ DNA, only 5% (11 of 216) of template locations had colocalized probe. Taken together, these results confirm that nearly all colocalization of probe with DNA template molecules indicates initiation at the *rrnB* P1 promoter on those molecules.

To determine the extent to which the UP element stimulates transcription from individual *rrnB* P1 DNA molecules, we next repeated the experiment using the DNA that lacks the UP element sequence (UP^−^; Fig. 1A). The mutation greatly reduced the frequency of initiation. To measure the rate of initiation on the DNA constructs under the conditions of the single molecule experiment, we measured the time to the first probe hybridization seen on each DNA (Fig. 2B). This time is the sum of the times required to initiate transcription on the DNA, to synthesize the first ∼45-68 nt of RNA needed to expose the hybridization targets, and to hybridize the probe. However, RNA synthesis (at ∼10 nt s^−1^ and hybridization; see Fig. S1) are >500-fold faster than initiation, so the time to the first hybridization is an accurate measure of the initiation time. Fits to the time distributions yielded the initiation rates (Fig. 2B); these confirmed that the UP^−^ construct retains some *rrnB* P1-dependent initiation activity. After subtracting the low rate of non-specific probe binding measured on UP^−^P1^−^, the P1-specific initiation rates at 0.75 nM RNAP on WT and UP^−^ were (3.7 ± 0.8) × 10^−4^ and (0.34 ± 0.10) × 10^−4^ s^−1^, respectively. Thus, elimination of the UP element from the wild-type promoter reduces by 11-fold the rate of transcription initiated from *rrnB* P1. This result is similar to that obtained in bulk experiments (Fig. S2) and agrees with the 11-fold decrease seen *in vivo* with the SUB-*lacZ* reporter construct [9].

### UP-Dependent Stimulation of Closed Complex Formation

The rate of initial association between σ^70^RNAP and promoters is thought to be a major step at which the rate of initiation is modulated by DNA sequence or transcription factors [5, 38, 39]. However, binding is most often measured in experiments which measure the initial association step only in aggregate with later steps [13]. To directly measure the second-order association rate, we observed binding of labeled σ^70^RNAP to surface-tethered DNA in the absence of NTPs [24, 25]. These experiments were conducted at >10-fold higher time resolution and excitation laser power relative to those in Fig. 2 to help ensure that even brief binding events were detected. Only the time to the first binding event on each DNA was scored in order to minimize any effect of occupancy of DNA by photobleached σ^70^RNAP. Distributions of times to first binding were exponential (Fig. 3A-D), yielding apparent first-order rate constants that were proportional to the σ^70^RNAP concentration (Fig. 3E). While there was little binding to areas of the chamber surface that did not contain DNA molecules, we observed significant binding to the UP^−^P1^−^ DNA molecules, presumably due to sequence-independent association of polymerase to the DNA or to binding of polymerase to non-P1 sequence elements (e.g., *rrnB* P2) present in the UP^−^P1^−^ construct.

**Figure 3.**
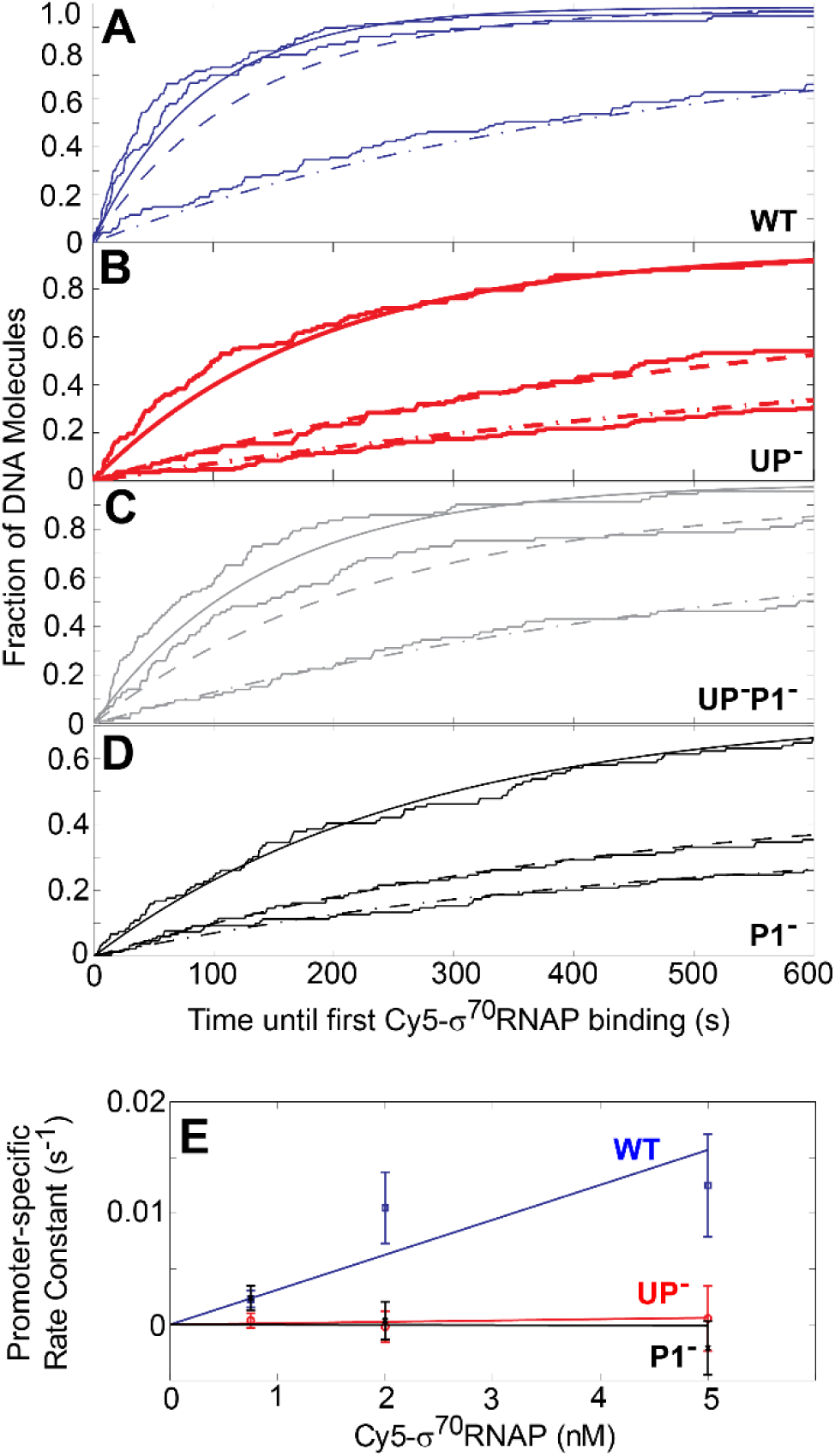
Effect of UP on association rate of Cy5-σ^70^RNAP to DNA. **(A-D)** Cumulative fraction of DNA molecules on which Cy5-σ^70^RNAP had appeared by the indicated time (data records) and exponential fits (smooth curves). Experiments were conducted in the absence of NTPs or probe with Cy5-σ^70^RNAP at 0.75 nM (dashed-dotted), 2 nM (dashed), and 5 nM (solid) on WT (A), UP^−^ (B), UP^−^P1^−^ (C) and P1^−^ (D) DNAs. Number of DNA molecules at each concentration was 85 – 166. (E) Apparent first-order rate constants (± SE; from fits in A-D) of promoter element-specific binding. In each case, the rate constant measured for UP^−^P1^−^ was subtracted from the rate constant measured for the indicated DNA to remove the effects of *rrnB* P1-independent binding, and the difference was corrected as described (see Methods). Proportional fits give second-order rate constants (3.1 ± 0.6) × 10^6^ M^−1^ s^−1^, (0.13 ± 0.4) × 10^6^ M^−1^ s^−1^, and (−0.01 ± 0.4) × 10^6^ M^−1^ s^−1^ for WT, UP^−^, and P1^−^ binding respectively. Imaging conditions: 0.5 s per frame; 1.25 mW 633 nm excitation.

The rate constants for association to the UP^−^ and P1^−^ constructs were indistinguishable within experimental uncertainty from the rate for UP^−^P1^−^. Thus, there was no detectable binding over background to the UP site alone or to the core promoter alone (Fig. 3E black and red). In contrast, the rate binding to WT DNA clearly exceeded binding to UP^−^P1^−^, yielding a second order rate constant for the promoter-specific binding of σ^70^RNAP to *rrnB* P1 of (3.1 ± 0.6) × 10^6^ M^−1^ s^−1^ (S.E.; Fig. 3E, blue). This value is >60-fold lower than a previous estimate of the lower limit on the association rate from filter binding experiments [9], but the difference is not unexpected given that the earlier work was done at lower ionic strength.

The estimated uncertainties in the measurements set upper limits (at *p* = 0.84) of <0.53 × 10^6^ and <0.39 × 10^6^ M^−1^ s^−1^ for specific binding of σ^70^RNAP to the P1-core and UP sites, respectively. However, the rate of binding at a given RNAP concentration must necessarily be greater than or equal to the rate of initiation at that concentration. Thus, the initiation rate measured above (Fig. 2) and the binding data constrain the P1-core specific binding rate to be in the range 0.045 × 10^6^ M^−1^ s^−1^ to 0.53 × 10^6^ M^−1^ s^−1^. Taken together, the data show that removal of UP decreases the rate of σ^70^RNAP binding to *rrnB* P1 by a factor that is at least 5.8-fold and may be as large as 69-fold.

### Promoter Complexes formed by σ^70^RNAP in the presence and absence of UP

Given the range of possible values, the increase in the binding rate of RNAP to *rrnB* P1 caused by UP accounts for most but possibly not all of the 11-fold increase in overall transcription rate caused by the presence of UP. Therefore, we next examined whether UP might also act at a subsequent step in initiation by altering the kinetic stabilities of holoenzyme-promoter complexes. To do this we examined the complex lifetimes in the absence of NTPs, conditions which allow formation of closed and open complexes but do not allow open complexes to proceed to RNA synthesis. Excitation power in this experiment was chosen to minimize photobleaching of Cy5-σ^70^RNAP (Fig. S3), in order to permit accurate measurement of the lifetimes of long-lived Cy5-σ^70^RNAP-DNA complexes. We observed that complex formation was highly reversible, with numerous short (< 0.5 min duration) holoenzyme binding events observed on nearly all WT DNAs (Fig. 4A, left). This behavior was qualitatively similar to that seen in the presence of NTPs in the experiments of Fig. 2. Conversely, we also detected some complexes that persisted for 5 min or more on WT DNA. UP^−^ displayed fewer such long lived complexes and the control UP^−^P1^−^ DNA almost none (Fig 4A). Thus, most observed complexes, whether of long or short lifetime, reflect sequence-specific promoter binding.

**Figure 4.**
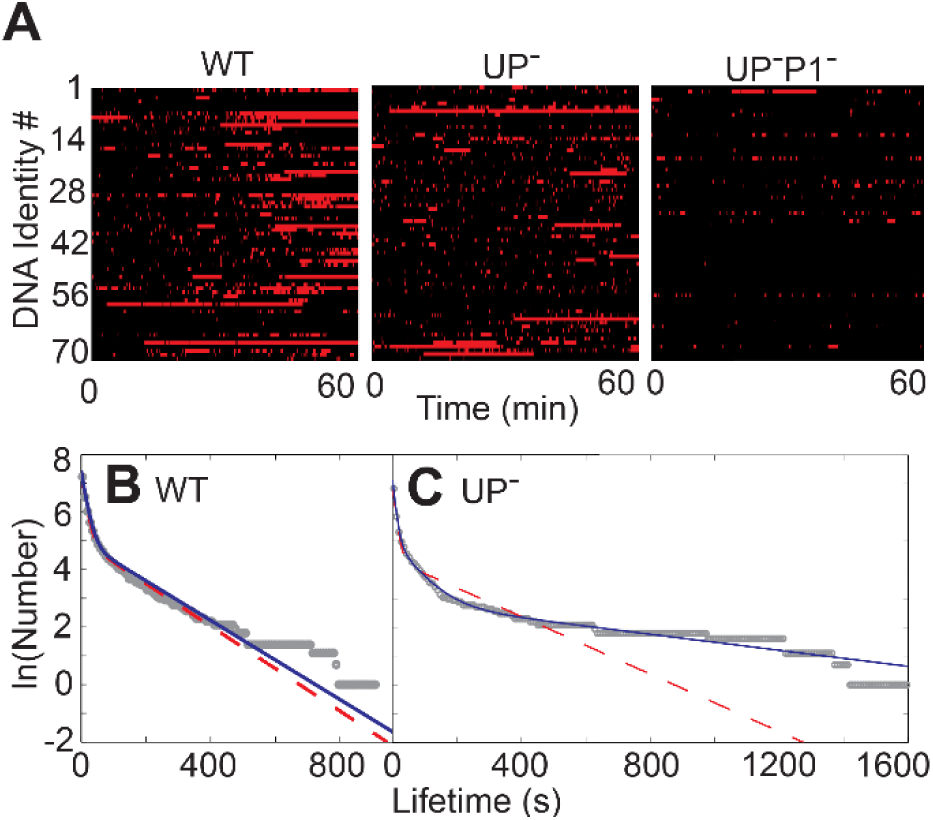
Effect of UP on σ^70^RNAP-DNA complex lifetimes. **(A)** Rastergrams illustrating Cy5-σ^70^RNAP binding events observed in separate individual experiments with WT, UP^−^, and UP^−^P1^−^ DNAs in the absence of NTPs. Imaging conditions: 8 s per frame; 25 µW 633 nm excitation. Each horizontal line in one of the rastergrams shows data from an individual DNA molecule (one of 72 randomly chosen from each experiment). Color indicates whether a colocalized spot of Cy5-σ^70^RNAP fluorescence was (red) or was not (black) present. **(B)** Cumulative distribution of *rrnB* P1-specific lifetimes (gray) of Cy5-σ^70^RNAP on WT DNA (*N* = 1,381). Each point represents the natural logarithm of the number of observed colocalization events with durations greater than or equal to the indicated lifetime. Graph also shows the distributions predicted from two-(red) and three-exponential (blue) fits to the lifetime data (see Methods and Table S2). **(C)** Same as B for UP^−^ DNA (*N* = 929). The three-exponential fit is decisively favored (ΔBIC_2−3_ = −10). In both (B) and (C) the frequency distribution of events observed on the UP^−^P1^−^ DNA has been subtracted from the observed frequencies to correct for the small amount of *rrnB* P1-independent binding.

Fitting the distribution of holoenzyme-WT DNA complex lifetimes required a function with at least two exponential terms (Fig. 4B). This is consistent with the known initiation mechanism: in the absence of NTPs, σ^70^RNAP can form at least two different complexes with *rrnB* P1 [9]. Unexpectedly, very long lived (> ∼500 s) complexes were observed more frequently on UP^−^ than on WT, although the number of such complexes was very small, representing only 1-2% of the total (Fig. 4C). This suggests that the absence of UP allows the formation of a small subpopulation of highly kinetically stable RNAP-promoter complexes that do not form when UP is present. Consistent with the presence of an additional species, statistical model selection decisively favored fitting the holoenzyme-UP^−^ DNA complex lifetime distribution with at least three exponential terms (Fig. 4C). Nevertheless, with the exception of rarely formed long-lived species on the UP^−^ DNA, the lifetime distributions on WT and UP^−^ DNA are similar, with roughly 90% of the lifetimes having a time constant of ∼10 s and the remainder ∼60 s or longer (Table S2).

### A Quantitative Kinetic Model for UP-Stimulated Transcription from WT *rrnB* P1

To gain further insight into the mechanistic implications of the data, we analyzed the results of the WT DNA binding and lifetime experiments in the context of a minimal sequential kinetic model in which an initial binding complex isomerizes into a second, more kinetically stable intermediate complex (Fig. 5A, black; we refer to the two complexes as RP_C_ and RP_I_, respectively). The model predicts exponential binding curves and bi-exponential lifetime distributions, corresponding to our observations (Fig. 3 and Fig. 4B, respectively). Values for rate constant *k*_1_ are defined as the promoter-specific binding rates (Fig. 3E). A two-exponential fit to the lifetime distribution of σ^70^RNAP on the WT DNA in the absence of NTPs (Fig. 4B) yielded estimates of the other rate constants (Fig. 5A, WT) for the initial steps of *rrnB* P1 initiation. In an otherwise identical lifetime distribution measurement conducted in the presence of ATP, CTP, and UTP, the amplitude and lifetime of the short component of the lifetime distribution were similar, while the long component lifetime(s) was significantly increased (Fig. S6A, Table S2). This increase is consistent with the expectation that NTPs do not affect the initial binding (step 1 in Fig. 5A) [13] but allow the formation of additional intermediates (TEC formed in the *k*_4_step) that are kinetically stabilized by initial RNA synthesis. If the post-isomerization steps (lumped together as the effective rate constant *k*_4_ in Fig. 5) do not limit the rate of initiation (which requires *k*_4_ >> *k*_-2_), the deduced rate constants derived from the model predict an initiation rate from WT *rrnB* P1 of (1.5 ± 0.4) × 10^−4^ s^−1^ at 0.75 nM RNAP This value is similar to the independently determined promoter-specific initiation rate (3.7 ± 0.8) × 10^−4^ s^−1^ (Fig. 2D), supporting the assumption that post-isomerization steps do not limit the rate. Thus, all of the WT DNA data are consistent with the minimal mechanism containing two kinetically significant complexes prior to the NTP-binding step, and the data support models that early reactions (up to and including the formation of the open complex) control the rate of initiation at *rrnB* P1.

**Figure 5.**
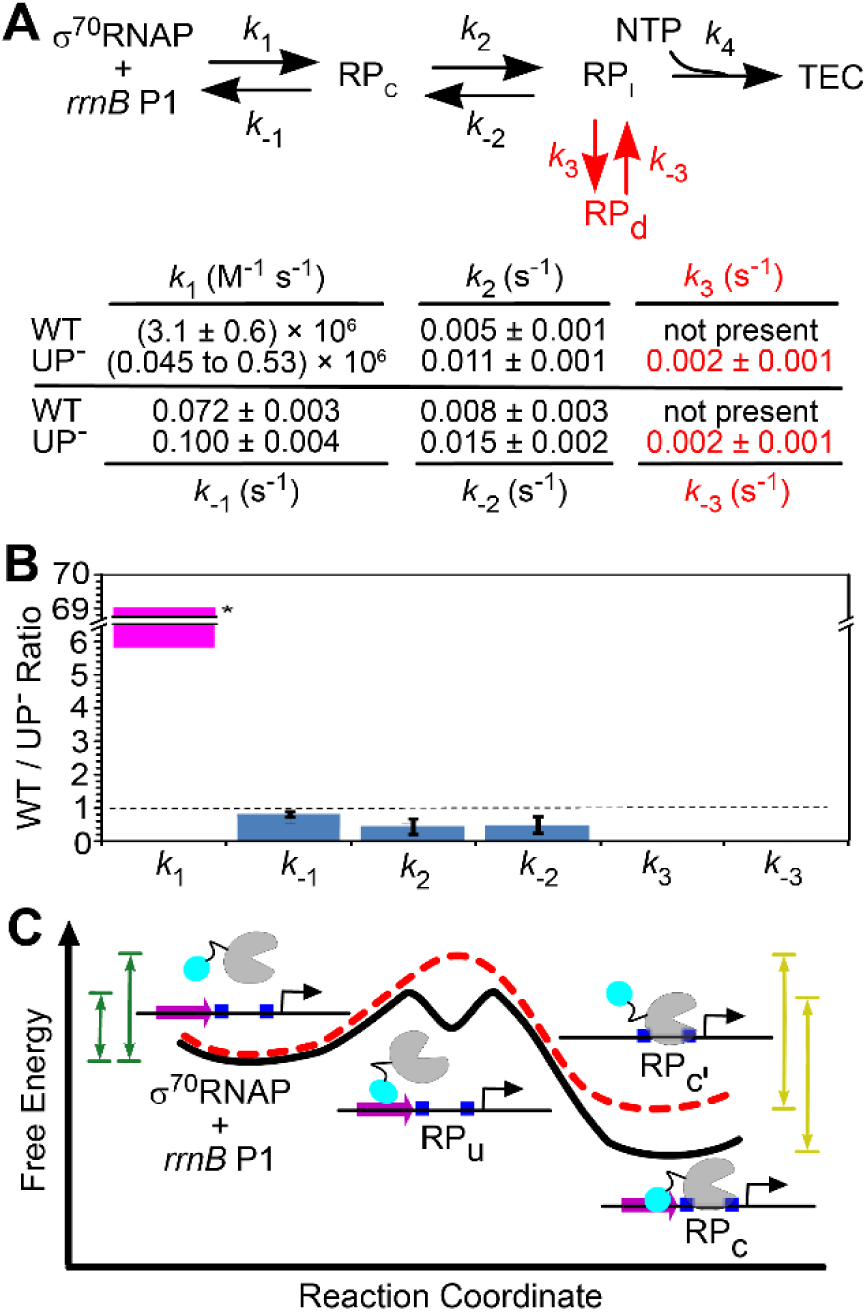
Effect of UP on the kinetic mechanism of *rrnB* P1 initiation. **(A)** Working minimal model of the reaction pathway for initiation and the values (±S.E) deduced from single-molecule data for the rate constants in the absence of NTPs. RP_c_ and RP_I_ are RNAP-promoter complexes that are intermediates on the initiation pathway; RP_d_ is an off-pathway state of unknown structure detected only on the UP^−^ DNA. RP_d_ may not be significantly populated except in the absence of NTPs (see text). Rate constant *k*_-3_ is the slowest first-order rate constant in the scheme and may be overestimated because of some contribution from photobleaching. **(B)** UP-induced fold change in the rate constants determined from fitting single molecule measurements with WT, UP^−^, and UP^−^P1^−^ DNAs to the working model. Asterisk marks value that is significantly greater than 1 (*p* < 0.01). **(C)** Postulated free energy landscapes for the initial binding step (*k*_1_ and *k*_-1_ in (A)) in the presence (black line) and absence (red line) of UP that explain the effects of UP (purple) and core promoter elements (blue) in σ^70^RNAP binding to and release from *rrnB* P1. The cyan circle represents the C-terminal domain of one of the α subunits of RNAP. Double-headed arrows indicate the free energy barriers that limit the rates of the association (pine) and dissociation (gold) reactions. Note that in the presence of UP, RP_C_ can also form from RP_Cꞌ_ (pathway not shown), but only a minority of molecules is expected to go through this pathway because of the higher barrier height on the red energy surface.

As previously discussed, lifetime distributions of σ^70^RNAP complexes on UP^−^ DNA require at least three exponential terms for a satisfactory fit even in the absence of NTPs (Fig. 4C). This implies the existence of three or more distinct species of RNAP-promoter complexes that can form prior to any step requiring NTPs. Thus, the two-intermediate model we used to interpret the WT DNA data (Fig. 5A, black only) is insufficient to explain the initiation mechanism on UP^−^ DNA. However, the three-intermediate model shown in Fig. 5A (black and red together) is consistent with the UP^−^ DNA lifetime distribution. Furthermore, the resulting rate constants predict a maximal initiation rate at 0.75 nM RNAP (assuming the upper limit value of *k*_1_, 0.53 × 10^6^ M^−1^ s^−1^, and that post-isomerization steps along the initiation pathway are fast) of (3.9 ± 0.3) × 10^−5^ s^−1^. This corresponds closely to the measured initiation rate on the UP^−^ promoter, (3.4 ± 1.0) × 10^−5^ s^−1^. Thus, the kinetic scheme shown in Fig. 5A is consistent with all of our experimental data on the UP^−^ DNA (Figs. 2B, 3E, and 4C).

The scheme of Fig. 5A includes a state, RP_d_, which is not a sequential intermediate in initiation but instead is positioned as a branch off the initiation pathway. The presence of this state would explain the small population (1.4% of binding events) of anonymously long-lived complexes that form on UP^−^ DNA in the absence of NTPs (Fig. 4C; Table S2). We considered an alternative model in which all three species were sequential intermediates along the initiation pathway (Fig. S7), but this model does not explain the rate of initiation on UP^−^ DNA, instead predicting a maximal initiation rate of (5.1 ± 2.4) × 10^−6^ s^−1^, ∼7-fold lower than that observed. Thus, it is unlikely that linear schemes with all intermediates on-pathway are sufficient to explain our observations. Despite the evidence for the off-pathway RP_d_ state under the artificial conditions of zero NTPs, this species is not necessarily important at physiological nucleotide concentrations. RP_d_ may not be significantly populated in the presence of NTPs if subsequent steps in initiation (*k*_4_ in Fig. 5A) are fast enough to effectively compete with its formation (*k*_3_).

## Discussion

We used single-molecule fluorescence to observe initiation, binding to and dissociation from rrnB P1 by single molecules of σ^70^RNAP. Unlike many previous studies of initiation, the experiments directly characterize the key initial promoter binding step. This reduces complications arising from coupling of this step to subsequent conformational isomerization of the RNAP-promoter complex [9, 13]. The experiments demonstrated that UP substantially stimulates (by 11-fold) initiation at this promoter *in vitro*. This value falls within the range measured for factor-independent UP stimulation of *rrnB* P1 *in vivo* [9]. The rate of sequence-dependent association of σ^70^RNAP with the core promoter was undetectably low but was substantially accelerated by addition of UP to the DNA construct. Reaction steps that interconvert the initial kinetically significant RNAP-DNA complexes were readily reversible both in the presence and absence of UP, and the rates of these steps and of polymerase dissociation from DNA were essentially unaffected by UP.

UP could potentially promote initiation either by increasing the rate of RNAP binding to the promoter or by increasing the efficiency with which bound complexes are converted to elongation complexes. The association measurements demonstrate that binding rate is increased by at least 6-fold, and possibly by much more since promoter sequence-specific binding without UP was too slow to detect in Fig. 3. The RNAP-DNA lifetime distributions show more subtle differences with and without UP, and the results of kinetic modeling suggest that initiation efficiency after initial binding is not increased by UP. Specifically, the initiation efficiency following promoter binding is calculated to be 6.5 ± 1.2% (S.E.) in the presence of UP and 9.9 ± 0.9% in its absence, subject to the assumption that post-isomerization steps are fast. Thus, UP does not significantly improve efficiency despite the fact that the presence of UP suppresses the production of a rare off-pathway state, RP_d_. Taken together, our data show that the stimulation of initiation by UP on *rrnB* P1 can be completely or nearly completely explained by stimulation of the recruitment of RNAP to the promoter in the initial binding step, at least at the sub-saturating concentrations of σ^70^RNAP used here.

### Comparison with previous studies of UP effects on initiation

There is extensive evidence, reviewed in [13], that DNA upstream of the −35 element can accelerate initiation by promoting isomerization to the open complex. However, some of these studies simply remove the upstream DNA and thus do not clearly distinguish between effects of the UP sequence from a generic involvement of upstream DNA in initiation step(s).

Other studies specifically examined the effects of mutation of UP sequences as we do here. Previous work that used bulk kinetic analyses based on filter binding methods [9] concluded that 1) UP stimulates the initial binding of σ^70^RNAP to *rrnB* P1, 2) UP stimulates (by ∼3-fold) isomerization of closed to open promoter complexes, and 3) UP blocks dissociation RNAP dissociation in that initial binding of RNAP to the promoter is reversible in the absence of UP but becomes effectively irreversible in its presence. Our data are compatible with the first but not with the second of these conclusions since we calculate detect little effect of UP on conversion efficiency indicating that any UP effect on closed-to-open complex isomerization likely plays a minimal role in UP stimulation of initiation. Our results disagree with the third conclusion: we directly observe that dissociation of the initial complex is fast (*k*_-1_ = ∼0.1 s^-1^). A possible reason for the discrepancies between our results and those of ref. [9] is that the former were conducted at monovalent cation concentrations that approximate those in live cells (150 mM KCl) [40] whereas the latter were conducted at lower salt concentrations that are known to reduce both dissociation and open complex formation [41, 42]. The method used here directly detects the initial binding step instead of monitoring it in aggregate with subsequent isomerizations and can do so even in physiological salt where binding is weak, conditions at which appropriate regulation of *rrnB* P1 by ppGpp and initiating nucleotide concentration [43] can be demonstrated *in vitro* [44].

### Mechanism by which UP accelerates binding of RNAP to promoter

The experimental results (Fig. 3E) demonstrate independently from any specific kinetic model that UP substantially stimulates the initial binding of polymerase to the promoter. We presume that the acceleration of *k*_1_ by UP (Fig. 5A,B) in the absence of upstream-binding factors like Fis [12] (ref) arise from initial binding of α-CTD to UP [13, 14], forming an intermediate complex RP_u_ along the pathway to RP_C_ (Fig. 5C). Since we detected (Fig. 3E) no binding of σ^70^RNAP above background levels to the P1^−^ DNA (which contains UP but lacks the −35 and −10 box promoter sequences), RP_U_ is likely to be kinetically unstable with a low barrier to dissociation. Nevertheless, its formation could significantly reduce the effective barrier to formation of RP_C_ as depicted in the free energy landscape in Fig. 5C, accelerating σ^70^RNAP binding to the promoter. This landscape is also consistent with the conclusion that dissociation of RP_C_ occurs at similar rates in the presence and absence of UP (*k*_-1_ in Fig. 5A, B), implying that the barriers to dissociation (gold arrows in Fig. 5C) are similar. This suggests that an UP-αCTD interaction stabilizes the closed complex (compare the free energies of RP_Cꞌ_ and RP_C_ in Fig. 5C), and is further evidence consistent with the involvement of the RP_u_ intermediate in initial σ^70^RNAP-DNA complex formation on the WT promoter.

How can the mere addition of a single additional binding target for RNAP accelerate the formation of RP_C_ by roughly an order of magnitude? UP is known to interact with the extreme C-terminal ends of the CTD domains of both α subunits of RNAP [11, 18, 45]. We speculate that the rate acceleration arises from the fact that the DNA-binding portion of α-CTD is a small domain [46, 47] on a long tether which is thought to be unstructured and highly flexible [45]. It is therefore expected to have greatly increased rotational and local translational Brownian motion relative to the much larger core polymerase domains. The high Brownian motion would in turn be expected to greatly increase the rate constant of the diffusion-controlled binding of the domain to its target site relative to the rate that the slowly diffusing core polymerase can achieve for binding to its target. A similar acceleration of the association rate constant by tethered diffusion of might also account for enhanced initiation by class I activators and other proteins which, like UP, serve as parts of upstream binding sites for α-CTD [1].

## Materials and Methods

### Construction of E. coli *rrnB* P1 template and its variants

Transcription templates were constructed by PCR using commercially obtained oligonucleotide primers (Table S1) labeled at their 5’-ends with either AlexaFluor 488 or biotin. PCR reactions used as template *E. coli* genomic DNA from strain MG1655 (ATCC), and high fidelity platinum Taq polymerase, and were run according to the manufacturer’s protocol (Stratagene). DNA sequences in the *rrnB* P1 positions were directed by the upstream PCR primer so that the entire promoter and UP element were present (WT), the UP element sequence was mutated (UP^−^), the −35 box of the P1 promoter was mutated (P1^−^), or both UP element and −35 box sequences were mutated (UP^−^P1^−^). All variants contained the genomic −27 to +240 sequence, including the weaker P2 promoter which is not the focus of this study, resulting in templates of 310 bp. For the UP^−^ and the UP^−^P1^−^ DNA variants, nucleotides −41 to −65 or −28 to −35 pyrimidines were changes to their non-pairing purine and purines were altered to their non-pairing pyrimidines respectively.

### Expression, Purification, and Labeling of *E. coli* σ^70^

Previously described and characterized [35] *rpoD*(Δcys) C132S C291S C295S S366C was cloned into pET15b for the purpose of adding an N-terminal His_6_-tag, yielding pRPODS366C (https://www.addgene.org/129688/). The plasmid was transformed into *E. coli* BL21DE3 pLysS, grown to OD_600_ = ∼0.4 at 37°C in 2× YT broth containing 100 µg/mL ampicillin, and induced by the addition of IPTG to 1 mM with further incubation for 2 hours. The expressed σ^70^S366C was purified according to previous protocols [48] with some minor changes: Cells were harvested and then separated from the periplasmic material by a series of buffer & osmotic strength changes [49] before dropwise freezing in liquid N_2_. Cells were thawed on ice in 50 mM Tris-Cl^−^ pH 8.0, 500 mM NaCl, 5 mM imidazole, 5% glycerol, 300 µg/mL lysozyme (Sigma-Aldrich) and complete protease inhibitor (Roche) before sonication. The soluble fraction of the cell lysate was then purified over a Talon nickel-chelating column as described previously [48]. σ^70^-containing fractions were then dialyzed against TGED buffer (50 mM Tris-Cl^−^ pH 8.0, 10% glycerol, 0.1 mM EDTA, and 0.1 mM DTT) to remove imidazole and further purified over a 5 mL HiTrapQ (GE Healthcare) anion exchange column. The protein eluted at 350 mM NaCl in a 50 mL, 100 – 500mM NaCl gradient in TGED buffer containing 50 μM TCEP instead of DTT. The addition of this anion-exchange chromatography step resulted in σ^70^ that was >95% pure determined by SDS-PAGE and Coomassie staining. Purified σ^70^S366C was mixed with a ten-fold molar excess of Cy5-maleimide (GE Healthcare) and incubated at room temperature for 10 min, followed by 2 hour incubation on ice. The resulting Cy5-σ^70^ was separated from excess dye on Sephadex G50 (0.6 × 28cm) at 4°C. Purified Cy5-σ^70^ was dialyzed against a storage buffer containing 10 mM Tris-Cl^−^ pH 8.0, 200 mM NaCl, 50% glycerol, 1mM EDTA, 1 mM DTT and stored at −20 °C. Removal of free dye from the labeled protein sample was confirmed by SDS-PAGE using a Typhoon scanner with a Cy5 filter set.

We determined the fraction labeled of Cy5-σ^70^ colorimetrically using ε_280_ = 39,040 M^−1^ cm^−1^ for σ^70^, and ε_650_ = 250,000 M^−1^ cm^−1^, ε_280_ / ε_650_ = 0.05 for Cy5. The results indicated that ∼70% of the protein was labeled with Cy5. The 30% dark σ^70^ was in rough agreement with the fraction of transcript probe binding events we observed in the absence of a preceding σ^70^-RNAP binding event.

### RNAP holoenzyme preparation

E. coli core RNAP (α_2_ββ′ω) with a SNAP-tag on the C-terminus of β′ [50] was a generous gift from Robert Landick, Rachel Mooney and Abbey Vangeloff and was labeled with SNAP-Surface 647 (New England Biolabs) as described [51] to produce core RNAP^647^.

To prepare σ^70^ holoenzyme, σ^70^ or Cy5-σ^70^ was removed from −20 °C storage and incubated with core RNAP or RNAP^647^ at a 1.3:1 mole ratio for ∼1 hr at 0 °C. Initiation rates for unlabeled σ^70^RNAP, Cy5-σ^70^RNAP, and σ^70^RNAP^647^ were identical within experimental uncertainty (Fig. S4). Four of six initiation rate data sets (two on WT DNA and two on UP^−^ DNA) were collected with σ^70^RNAP^647^; all other data were collected with Cy5-σ^70^RNAP.

### Flow chamber preparation and microscopy

The micro-mirror multiwavelength single-molecule total internal reflection (TIR) fluorescence microscope used excitation lasers at 488, 532, and 633 nm and was previously described [24]. Reactions were performed in ∼20 μl flow chambers (∼4 × ∼25 × ∼0.2 mm). Labeled DNA surface density was ∼0.15 molecules μm^−2^, yielding a template concentration averaged over the cell volume of ∼1.2 pM. Biotinated, dye labeled DNA molecules were linked with avidin DN (Vector Laboratories) to the surface of a fused silica flow cell essentially as described [24]. Reactions with σ^70^RNAP were performed in microscopy buffer: 40 mM Tris-OAc (pH 8.0), 150 mM KCl, 10 mM MgCl_2_, 4 mM DTT, 10 mg/ml bovine serum albumin (#126615 EMD Chemicals; La Jolla, CA), and a PCA/PCD oxygen scavenging system [52] to minimize photobleaching. Unless otherwise specified, reactions contained 0.75 nM σ^70^RNAP. Transcription reactions were conducted in microscopy buffer supplemented with 0.5 mM of each NTP and 2.5 nM of each Cy3 oligonucleotide probe.

### Analysis of CoSMoS Data

Analysis of single-molecule colocalization data was performed with custom software implemented in Matlab as previously described [24, 53] with minor changes. To score association and release of labeled molecules (σ^70^RNAP or transcript probe oligonucleotide) from a surface-tethered DNA molecule, we integrated the fluorescence emission of the molecules over a 0.45 × 0.45 µm areas centered at DNA locations. In the resulting records, increases to an emission intensity > 4.0 times the standard deviation of the baseline noise were scored as a binding event. Following a binding event, the first decrease of the emission to < 1.5 times the standard deviation of the baseline noise was scored as dissociation/photobleaching.

### Determination of σ^70^-RNAP initiation rates from *E. coli rrnB* P1

The rates and active fractions from three initiation rate data sets on WT DNA were determined by maximum likelihood fitting of the times to first observation of the transcript probe fluorescence on each DNA molecule and of the number of DNA molecules that showed no transcript signal throughout the duration of the experimental record [53]. The likelihood function assumed an exponential probability density. Standard errors of the rate and initiation fraction fit parameters were estimated by bootstrapping (random sampling, with replacement, from the full set of DNA molecules observed, which includes both molecules that did and those that did not initiate). The final rate [(3.8 ± 0.8) × 10^−4^ s^−1^] and active fraction [0.99 ± 0.02] values were determined by weighted averaging of the fit parameters from the three data sets. Data from experiments on the UP^−^ and UP^−^P1^−^ DNAs was processed the same way, except that 1) the individual data sets (three for UP^−^ and two for UP^−^P1^−^) were pooled and fit as a group (this was possible because all data sets had the same duration), and 2) the active fraction parameter was not fit but instead was fixed at 0.99, the value determined from the WT data (this was necessary because the slow rates of initiation of these DNA prevented accurate determination of the active fraction on the time scale observed).

### Determination of σ^70^-RNAP binding rates

To measure binding rates, microscopy buffer containing Cy5-σ^70^RNAP at 0.75 nM, 2 nM, or 5 nM was introduced into the flow chamber at time zero. Cy5-σ^70^RNAP was excited with 1.25 mW 633 nm light and fluorescence emission >635 nm was recorded at a frame rate of 2 Hz with autofocus adjustment every 240 frames. Under this high excitation power conditions, short binding events were efficiently detected but lasted only a few frames, presumably due to photobleaching. Binding events that colocalized with surface DNA positions were scored using an automated spot-picking algorithm [53]. Only the interval between time zero and the first σ^70^RNAP colocalization event on each DNA was scored; all subsequent binding events were ignored to minimize the effect of DNA occupancy by photobleached complexes on the measured rates. DNA molecules already bound with holoenzyme at time zero were excluded from the analysis.

For each set of measurements, an apparent first-order rate constant was determined by maximizing the single exponential likelihood function (Fig. 3A-D). The rate constants for specific binding to the promoter or its constituent elements (“promoter element-specific” rate constants) were determined by subtracting the apparent first-order rate constant measured with the UP^−^P1^−^ DNA from those measured with the WT, UP^−^, or P1^−^ DNAs at the same polymerase concentration (Fig. 3E). The specific binding rate constants were then adjusted by applying two correction factors to yield the final value of the rate constant. First, the rate constant was multiplied by 1.4 to account for the labeling stoichiometry of σ^70^RNAP measured in bulk. Second, the rate constant was multiplied by a factor to account for the binding events that were of durations too short (typically <0.4 s) to be experimentally detected due to background noise. To determine the size of this correction factor, we first selected in a typical recording every fluorescent spot with a duration ≥3 frames. In each frame in which the spot was visible, except the first and last, we measured the spot width and peak fluorescence intensity above background by Gaussian fitting (Fig. S5A). Next, we generated simulated spot images. Each image was constructed by adding a Gaussian function of the same width as the median width of the experimentally measured spots and a chosen peak intensity to a background noise image taken from a randomly selected frame and position in a spot-free region of an experimental record. For each Gaussian peak intensity value, we determined the maximum intensity threshold at which the simulated spot was detected by the spot-picking algorithm (Fig. S5B). The shortest duration peak spot that could be detected in the experimental records was then estimated to be

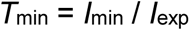

where *I*_min_ was the minimum Gaussian peak intensity detected at the threshold used in the analysis of the experimental data (46.2; Fig. S5B dashed lines) and *I*_exp_ was the median intensity measured for spots in the experimental images (Figure S5A, horizontal dashed line). This analysis was repeated once on each day data was recorded. Finally, the fraction of events that were too short to be detected was calculated as the integral of the experimental spot lifetime distribution from zero to *T*_min_. The rate constant was then multiplied by the reciprocal of this faction (which ranged from 1.2 to 1.4) to yield the final value.

### Determination of σ^70^RNAP-DNA complex lifetimes

To determine complex lifetimes, labeled σ^70^RNAP was diluted into microscopy buffer (final concentration 0.75 nM) and introduced into the flow chamber. σ^70^RNAP fluorescence was excited by 25 µW 633 nm light and emission >635 nm was recorded at 0.13 Hz with automatic focus adjustment [54] every 17 frames. The low frame rate reduced the time resolution of the experiment relative to that in the binding rate measurements, but allowed excitation at greatly reduced power, which minimized photobleaching. The duration that each polymerase molecule stayed on the DNA was scored by the threshold-crossing algorithm used previously [24], with the onset of binding scored when the intensity first exceeded 5.0 times the baseline noise and end of binding scored when the intensity dropped below 3.0 times the baseline noise. This procedure efficiently detected binding durations as short as ∼3 s. All binding events were inspected manually and those that did not image as a well-defined fluorescent spot were discarded. Dwell times were first measured in control experiments using UP^−^P1^−^ DNA. Since this DNA lacks the core *rrnB* P1 elements, DNA-colocalized binding events scored in these experiments result from *rrnB* P1-independent events such as σ^70^RNAP binding to the flow cell surface near a DNA molecule, to non-promoter regions of the DNA, or to *rrnB* P2. All of the binding events observed in this control were of short duration: most (>95%) lasted for one 8 s frame and the remainder lasted either 2 or 3 frames. The frequencies of these *rrnB* P1-independent events were subtracted from the frequencies of the 1-3 frame duration events measured with the WT, or UP^−^ DNAs to determine the distributions *of rrnB* P1-specific events [24]. Measured dwell times were fit to a two- or three-exponential lifetime distribution truncated at *t*_min_ = 3s. For WT DNA 1,381 of an estimated total of 1,433 binding events were detected; for UP^−^ 929 of an estimated 1,248 events were detected.

### Calculation of rate constants for formation and breakdown of σ^70^RNAP-promoter complexes

Rate constants *k*_-1_, *k*_2_, and *k*_-2_ for WT DNA and *k*_-1_, *k*_2_, *k*_-2_, *k*_3_, and *k*_-3_ for UP^−^ DNA (Fig. 5A) were calculated from the two- or three-exponential lifetime distribution fit parameters (Table S2) by iterative optimization using the Q-matrix method [55]. We assumed that *k*_4_ = 0 in the absence of nucleotide. The Bayesian information criterion (BIC) used for model selection was calculated as described [56]. Standard errors of the rate constants were estimated by bootstrapping [24]. The efficiency of initiation from RP_C_ in the model of Fig. 5A was calculated (based on the assumption that *k*_4_ >> [*k*_3_ + *k_−_*_2_]) as *k*_2_ / (*k*_-1_ + *k*_2_) and used the rate constants from the individual bootstrap trials to propagate the error estimates. In the alternate model (Fig. S7) the efficiency of initiation from RP_C_ was calculated as *k*_2_ *k*_3_ / [*k*_−1_ (*k*_−2_ + *k*_3_) + *k*_2_ *k*_3_], based on the assumption that *k*_4_ >> *k*_-3_.

### Transcript probe hybridization rate

To measure the rate constant for transcript probe hybridization, we first pre-formed TEC complexes stalled at the avidin-biotin roadblock by incubating surface-anchored DNA with σ^70^RNAP and NTPs. Following a one hour of incubation, buffer containing 2.5 nM each Cy3-transcript probe was introduced and the appearance over time of Cy3 fluorescence spots localizing with transcript in the stalled TECs was recorded. The distribution of times for the appearance of the first hybridization at each location was fit to a single exponential, yielding a mean transcript detection time of 35 s.

## Acknowledgements

We thank R. Mooney, R. Landick, B. Smith, and M. Marr for providing materials and for helpful discussions. This work was supported by grant R01GM81648 from NIGMS.

## References

1. Browning DF, Busby SJW (2016) Local and global regulation of transcription initiation in bacteria. Nature Reviews Microbiology, 14:638–650. https://doi.org/10.1038/nrmicro.2016.103

2. Werner F, Grohmann D (2011) Evolution of multisubunit RNA polymerases in the three domains of life. Nature Reviews. Microbiology, 9(2):85–98. https://doi.org/10.1038/nrmicro2507

3. Sainsbury S, Bernecky C, Cramer P (2015) Structural basis of transcription initiation by RNA polymerase II. Nature Reviews Molecular Cell Biology, 16(3):129–143. https://doi.org/10.1038/nrm3952

4. Mosaei H, Harbottle J (2019) Mechanisms of antibiotics inhibiting bacterial RNA polymerase. Biochemical Society Transactions, 47(1):339–350. https://doi.org/10.1042/BST20180499

5. Minchin SD, Busby SJW (2009) Analysis of mechanisms of activation and repression at bacterial promoters. Methods (San Diego, Calif.), 47(1):6–12. https://doi.org/10.1016/j.ymeth.2008.10.012

6. Bartlett MS, Gaal T, Ross W, Gourse RL (2000) Regulation of rRNA transcription is remarkably robust: FIS compensates for altered nucleoside triphosphate sensing by mutant RNA polymerases at Escherichia coli rrn P1 promoters. Journal of Bacteriology, 182(7):1969. https://doi.org/10.1128/jb.182.7.1969-1977.2000

7. Dickson R, Gaal T, deBoer H, deHaseth PL, Gourse RL (1989) Identification of promoter mutants defective in growth rate-dependent regulation of rRNA transcription in Escherichia coli. Journal of Bacteriology, 171:4862–4870. https://doi.org/10.1128/jb.171.9.4862-4870.1989

8. Bartlett MS, Gourse RL (1994) Growth rate-dependent control of the rrnB P1 core promoter in Escherichia coli. Journal of Bacteriology, 176(17):5560–5564. https://doi.org/10.1128/jb.176.17.5560-5564.1994

9. Rao L, Ross W, Appleman JA, Gaal T, Leirmo S, Schlax PJ, Record MT Jr, Gourse RL (1994) Factor independent activation of rrnB P1. An “extended” promoter with an upstream element that dramatically increases promoter strength. Journal of Molecular Biology, 235(5):1421–1435. https://doi.org/10.1006/jmbi.1994.1098

10. Newlands JT, Ross W, Gosink KK, Gourse RL (1991) Factor-independent activation of Escherichia coli rRNA transcription II. Characterization of complexes of rrnB P1 promoters containing or lacking the upstream activator region with Escherichia coli RNA polymerase. Journal of Molecular Biology, 220(3):569–583.

11. Estrem ST, Ross W, Gaal T, Chen ZW, Niu W, Ebright RH, Gourse RL (1999) Bacterial promoter architecture: subsite structure of UP elements and interactions with the carboxy-terminal domain of the RNA polymerase alpha subunit. Genes & Development, 13(16):2134– 2147. https://doi.org/10.1101/gad.13.16.2134

12. Hirvonen CA, Ross W, Wozniak CE, Marasco E, Anthony JR, Aiyar SE, Newburn VH, Gourse RL (2001) Contributions of UP Elements and the Transcription Factor FIS to Expression from the Seven rrn P1 Promoters in Escherichia coli. Journal of Bacteriology, 183(21):6305– 6314. https://doi.org/10.1128/JB.183.21.6305-6314.2001

13. Saecker RM, Record MT Jr, Dehaseth PL (2011) Mechanism of bacterial transcription initiation: RNA polymerase - promoter binding, isomerization to initiation-competent open complexes, and initiation of RNA synthesis. Journal of Molecular Biology, 412(5):754–771. https://doi.org/10.1016/j.jmb.2011.01.018

14. Borukhov S, Nudler E (2008) RNA polymerase: the vehicle of transcription. Trends in Microbiology, 16(3):126–134. https://doi.org/10.1016/j.tim.2007.12.006

15. Cellai S, Mangiarotti L, Vannini N, Naryshkin N, Kortkhonjia E, Ebright RH, Rivetti C (2007) Upstream promoter sequences and αCTD mediate stable DNA wrapping within the RNA polymerase–promoter open complex. EMBO reports, 8(3):271–278. https://doi.org/10.1038/sj.embor.7400888

16. Chen H, Tang H, Ebright RH (2003) Functional interaction between RNA polymerase alpha subunit C-terminal domain and sigma70 in UP-element- and activator-dependent transcription. Molecular Cell, 11(6):1621–1633. https://doi.org/10.1016/s1097-2765(03)00201-6

17. Dove SL, Huang FW, Hochschild A (2000) Mechanism for a transcriptional activator that works at the isomerization step. Proceedings of the National Academy of Sciences of the United States of America, 97(24):13215–13220. https://doi.org/10.1073/pnas.97.24.13215

18. Ross W, Gourse RL (2005) Sequence-independent upstream DNA-alphaCTD interactions strongly stimulate Escherichia coli RNA polymerase-lacUV5 promoter association. Proceedings of the National Academy of Sciences of the United States of America, 102(2):291–296. https://doi.org/10.1073/pnas.0405814102

19. Ohlsen KL, Gralla JD (1992) Melting during steady-state transcription of the rrnB P1 promoter in vivo and in vitro. Journal of Bacteriology, 174(19):6071–6075. https://doi.org/10.1128/jb.174.19.6071-6075.1992

20. Sander P, Langert W, Mueller K (1993) Mechanisms of upstream activation of the rrnD promoter P1 of Escherichia coli. The Journal of Biological Chemistry, 268(23):16907–16916.

21. Strainic MG, Sullivan JJ, Velevis A, deHaseth PL (1998) Promoter Recognition by Escherichia coli RNA Polymerase: Effects of the UP Element on Open Complex Formation and Promoter Clearance. Biochemistry, 37(51):18074–18080. https://doi.org/10.1021/bi9813431

22. Doniselli N, Rodriguez-Aliaga P, Amidani D, Bardales JA, Bustamante C, Guerra DG, Rivetti C (2015) New insights into the regulatory mechanisms of ppGpp and DksA on Escherichia coli RNA polymerase-promoter complex. Nucleic Acids Research, 43(10):5249–5262. https://doi.org/10.1093/nar/gkv391

23. Sreenivasan R, Heitkamp S, Chhabra M, Saecker R, Lingeman E, Poulos M, McCaslin D, Capp MW, Artsimovitch I, Record MT (2016) Fluorescence Resonance Energy Transfer Characterization of DNA Wrapping in Closed and Open *Escherichia coli* RNA Polymerase−λP R Promoter Complexes. Biochemistry, 55(14):2174–2186. https://doi.org/10.1021/acs.biochem.6b00125

24. Friedman LJ, Gelles J (2012) Mechanism of Transcription Initiation at an Activator-Dependent Promoter Defined by Single-Molecule Observation. Cell, 148(4):679–689. https://doi.org/10.1016/j.cell.2012.01.018

25. Friedman LJ, Mumm JP, Gelles J (2013) RNA polymerase approaches its promoter without long-range sliding along DNA. Proceedings of the National Academy of Sciences of the United States of America, 110(24):9740–9745. https://doi.org/10.1073/pnas.1300221110

26. Stumper SK, Ravi H, Friedman LJ, Mooney RA, Corrêa IR, Gershenson A, Landick R, Gelles J (2019) Delayed inhibition mechanism for secondary channel factor regulation of ribosomal RNA transcription. eLife, 8:e40576. https://doi.org/doi:10.7554/eLife.40576

27. Revyakin A, Allemand JF, Croquette V, Ebright RH, Strick TR (2003) Single-molecule DNA nanomanipulation: detection of promoter-unwinding events by RNA polymerase. Methods in Enzymology, 370:577–598. https://doi.org/10.1016/S0076-6879(03)70049-4

28. Larson MH, Landick R, Block SM (2011) Single-Molecule Studies of RNA Polymerase: One Singular Sensation, Every Little Step It Takes. Molecular Cell, 41(3):249–262. https://doi.org/10.1016/j.molcel.2011.01.008

29. Dangkulwanich M, Ishibashi T, Bintu L, Bustamante C (2014) Molecular Mechanisms of Transcription through Single-Molecule Experiments. Chemical Reviews, 114(6):3203–3223. https://doi.org/10.1021/cr400730x

30. Murray HD, Appleman JA, Gourse RL (2003) Regulation of the Escherichia coli rrnB P2 promoter. Journal of Bacteriology, 185(1):28. https://doi.org/10.1128/jb.185.1.28-34.2003

31. Friedman LJ, Chung J, Gelles J (2006) Viewing dynamic assembly of molecular complexes by multi-wavelength single-molecule fluorescence. Biophysical Journal, 91(3):1023–1031. https://doi.org/10.1529/biophysj.106.084004

32. Hoskins AA, Friedman LJ, Gallagher SS, Crawford DJ, Anderson EG, Wombacher R, Ramirez N, Cornish VW, Gelles J, Moore MJ (2011) Ordered and Dynamic Assembly of Single Spliceosomes. Science, 331(6022):1289–1295. https://doi.org/10.1126/science.1198830

33. Harden TT, Wells CD, Friedman LJ, Landick R, Hochschild A, Kondev J, Gelles J (2016) Bacterial RNA polymerase can retain σ70 throughout transcription. PNAS, 113:602–7. https://doi.org/10.1073/pnas.1513899113

34. Harden TT, Herlambang KS, Chamberlain M, Lalanne J-B, Wells CD, Li G-W, Landick R, Hochschild A, Kondev J, Gelles J (2020) Alternative transcription cycle for bacterial RNA polymerase. Nature Communications, 11(1):448. https://doi.org/10.1038/s41467-019-14208-9

35. Callaci S, Heyduk E, Heyduk T (1998) Conformational Changes of Escherichia coli RNA Polymerase σ70 Factor Induced by Binding to the Core Enzyme. Journal of Biological Chemistry, 273(49):32995. https://doi.org/10.1074/jbc.273.49.32995

36. Perlow RA (2003) Construction and purification of site-specifically modified DNA templates for transcription assays. Nucleic Acids Research, 31(7):40e–440. https://doi.org/10.1093/nar/gng040

37. King RA, Sen R, Weisberg RA (2003) Using a lac Repressor Roadblock to Analyze the E. Coli Transcription Elongation Complex. Methods in Enzymology, 371:207–218. https://doi.org/10.1016/S0076-6879(03)71015-5

38. Browning DF, Busby SJ (2004) The regulation of bacterial transcription initiation. Nature Reviews Microbiology, 2(1):57–65. https://doi.org/10.1038/nrmicro787

39. Roy S, Garges S, Adhya S (1998) Activation and repression of transcription by differential contact: two sides of a coin. The Journal of Biological Chemistry, 273(23):14059–14062. https://doi.org/10.1074/jbc.273.23.14059

40. Cayley S, Lewis BA, Guttman HJ, Record MT (1991) Characterization of the cytoplasm of Escherichia coli K-12 as a function of external osmolarity. Journal of Molecular Biology, 222(2):281–300. https://doi.org/10.1016/0022-2836(91)90212-O

41. Gourse RL (1988) Visualization and quantitative analysis of complex formation between E. coli RNA polymerase and an rRNA promoter in vitro. Nucleic Acids Research, 16(20):9789– 9809. https://doi.org/10.1093/nar/16.20.9789

42. Ohlsen KL, Gralla JD (1992) Interrelated effects of DNA supercoiling, ppGpp, and low salt on melting within the Escherichia coli ribosomal RNA rrnB P1 promoter. Molecular Microbiology, 6(16):2243–2251. https://doi.org/10.1111/j.1365-2958.1992.tb01400.x

43. Paul BJ, Barker MM, Ross W, Schneider DA, Webb C, Foster JW, Gourse RL (2004) DksA: A Critical Component of the Transcription Initiation Machinery that Potentiates the Regulation of rRNA Promoters by ppGpp and the Initiating NTP. Cell, 118(3):311–322. https://doi.org/10.1016/j.cell.2004.07.009

44. Barker MM, Gaal T, Josaitis CA, Gourse RL (2001) Mechanism of regulation of transcription initiation by ppGpp. I. Effects of ppGpp on transcription initiation in vivo and in vitro. Journal of Molecular Biology, 305(4):673–688. https://doi.org/10.1006/jmbi.2000.4327

45. Blatter EE, Ross W, Tang H, Gourse RL, Ebright RH (1994) Domain organization of RNA polymerase alpha subunit: C-terminal 85 amino acids constitute a domain capable of dimerization and DNA binding. Cell, 78(5):889–896. https://doi.org/10.1016/s0092-8674(94)90682-3

46. Benoff B, Yang H, Lawson CL, Parkinson G, Liu J, Blatter E, Ebright YW, Berman HM, Ebright RH (2002) Structural Basis of Transcription Activation: The CAP-αCTD-DNA Complex. Science, 297(5586):1562–1566. https://doi.org/10.1126/science.1076376

47. Liu B, Hong C, Huang RK, Yu Z, Steitz TA (2017) Structural basis of bacterial transcription activation. Science, 358(6365):947–951. https://doi.org/10.1126/science.aao1923

48. Marr MT, Roberts JW (1997) Promoter recognition as measured by binding of polymerase to nontemplate strand oligonucleotide. Science, 276(5316):1258–1260. https://doi.org/10.1126/science.276.5316.1258

49. Magnusdottir A, Johansson I, Dahlgren L-G, Nordlund P, Berglund H (2009) Enabling IMAC purification of low abundance recombinant proteins from E. coli lysates. Nature Methods, 6(7):477–478. https://doi.org/10.1038/nmeth0709-477

50. Tetone LE, Friedman LJ, Osborne ML, Ravi H, Kyzer S, Stumper SK, Mooney RA, Landick R, Gelles J (2017) Dynamics of GreB-RNA polymerase interaction allow a proofreading accessory protein to patrol for transcription complexes needing rescue. Proceedings of the National Academy of Sciences, 114(7):E1081–E1090. https://doi.org/10.1073/pnas.1616525114

51. Sanchez A, Osborne ML, Friedman LJ, Kondev J, Gelles J (2011) Mechanism of transcriptional repression at a bacterial promoter by analysis of single molecules. The EMBO Journal, 30:3940–3946. https://doi.org/10.1038/emboj.2011.273

52. Patil PV, Ballou DP (2000) The use of protocatechuate dioxygenase for maintaining anaerobic conditions in biochemical experiments. Analytical Biochemistry, 286(2):187–192. https://doi.org/10.1006/abio.2000.4802

53. Friedman LJ, Gelles J (2015) Multi-wavelength single-molecule fluorescence analysis of transcription mechanisms. Methods, 86:27–36. https://doi.org/10.1016/j.ymeth.2015.05.026

54. Crawford DJ, Hoskins AA, Friedman LJ, Gelles J, Moore MJ (2007) Visualizing the splicing of single pre-mRNA molecules in whole cell extract. RNA, 14(1):170–179. https://doi.org/10.1261/rna.794808

55. Colquhoun, D, Hawkes, AG (1995) A Q-Matrix Cookbook. Single-Channel Recording, :589–633.

56. Horn R (1987) Statistical methods for model discrimination. Applications to gating kinetics and permeation of the acetylcholine receptor channel. Biophysical Journal, 51(2):255–263. https://doi.org/10.1016/S0006-3495(87)83331-3

