## Supplemental figures and tables for "Mechanism of upstream promoter element stimulation of transcription at a ribosomal RNA promoter determined by single-molecule imaging"

Supplemental Figures S1 through S7

Supplemental Tables S1 and S2

### Supplemental Figures

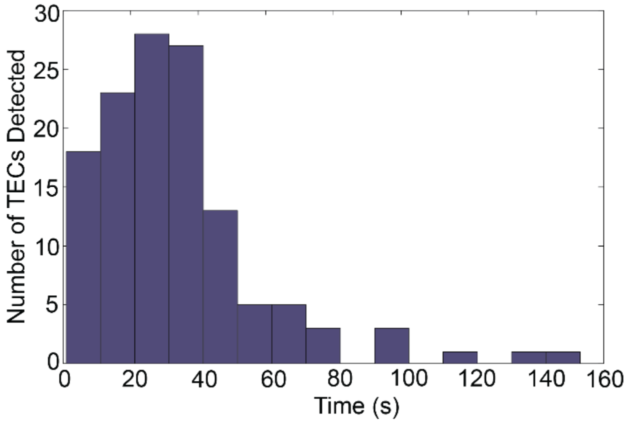


**Figure S1. Distribution of the time to detect the transcript by hybridization of one of the transcript probes to pre-formed TECs.** TECs roadblocked at +251 were pre-formed by 60 min incubation of surface-tethered WT DNA with β′-labeled $\sigma^{70}$RNAP and 0.5 mM NTPs and detected by the presence of the RNAP fluorescence. At time zero, the two Cy3-labeled transcript probes (2.5 nM each) were introduced and the time at which a probe was detected at each TEC location was determined using 500 μW 532 nm excitation at a frame rate of 0.033 Hz with an interruption every 17 frames for automatic focusing. The mean of the distribution shown is 33.4 s.

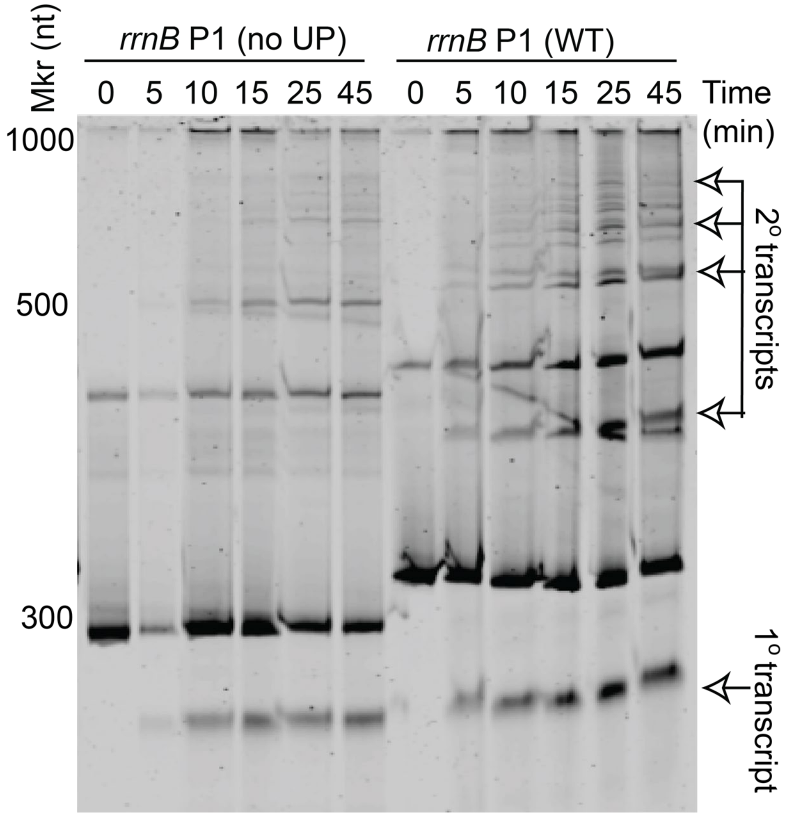


**Figure S2. Denaturing gel electrophoresis of run-off transcripts produced from WT DNA and from a variant in which UP is removed by truncation at -41.** Reactions were performed on 10 nM DNA in 1 mM NTPs and initiated at time zero by addition of 100 nM $\sigma^{70}$RNAP at 25°C. At the indicated times, aliquots were quenched by the addition of EDTA to a final concentration of 20 mM, extracted from an equal volume of 25:24:1 (v/v/v) phenol:chloroform:isoamyl alcohol and run on a 5.5% polyacrylamide gel containing 8M urea. The gel was stained for 20 min in a 1:10,000 dilution of SYBR Green II (Invitrogen) and imaged using a Typhoon scanner (GE Healthcare) with 488 nm excitation and a Cy3 filter set with photomultiplier at 380V. Primary and secondary transcript bands are indicated.

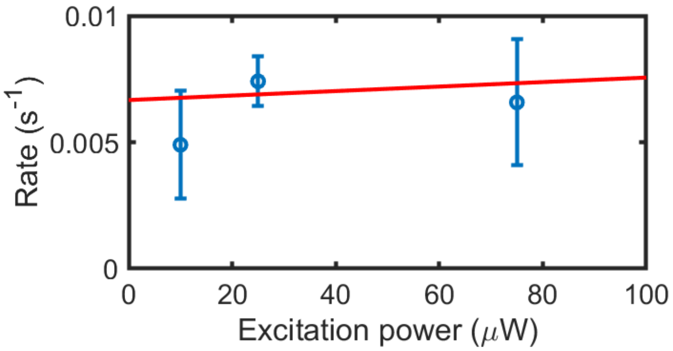


**Figure S3. Cy5-σ$^{70}$RNAP photobleaching kinetics.** Disappearance rates (blue; ±SE.) corresponding to the long-lifetime component of Cy5-σ$^{70}$RNAP bound to WT DNA in the absence of NTPs at 633 nm excitation powers of 10, 25, or 75 μW were calculated as 1 / $\tau_2$ (as in Fig. 4B and Table S2). Slope of the linear fit (red) was not significantly different from zero (95% confidence interval: [−6.3, 6.5] $\times$ $10^{-4}$ $s^{-1}$ $\mu W^{-1}$), demonstrating that there was no experimentally detectable dependence of the rate on laser power.

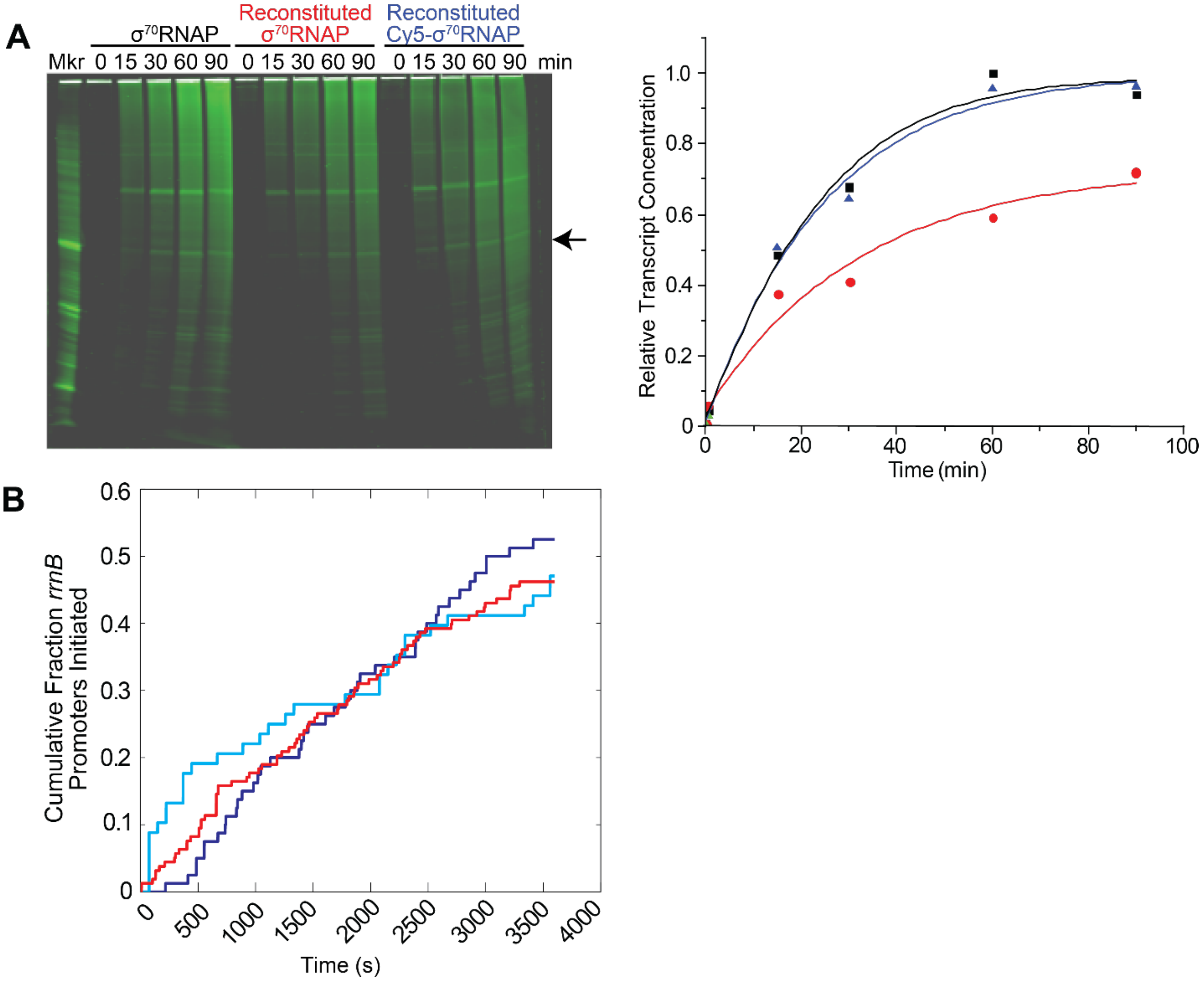


**Figure S4. Initiation activity on *rrnB* P1 of labeled and unlabeled σ$^{70}$RNAP preparations. (A)** Bulk *in vitro* transcription assay using 100 nM σ$^{70}$RNAP holoenzyme (black, Epicentre), 100 nM core RNAP (Epicentre) mixed with 500 nM wild type σ$^{70}$ (red), or 100 nM core RNAP mixed with 500 nM Cy5-σ$^{70}$ (blue). Reactions containing 15 nM T7A1 promoter template, and 750 μM NTPs were quenched after 30°C incubation at the indicated times by adding EDTA to a final concentration of 20 mM. The 6% polyacrylamide gel containing 8 M urea was stained for 20 min with a 1:10,000 dilution of SYBR Green II and scanned with a Typhoon scanner using 488 nm excitation (380V) and a Cy3 filter set (left). Relative amounts of transcript produced over time (right) were determined by integration of the full-length product band (arrow). Mkr, marker. **(B)** Single molecule initiation measurements (as in Fig. 2B) on WT *rrnB* P1 DNA using Cy5-σ$^{70}$RNAP (blue) or σ$^{70}$RNAP$^{647}$ (two replicates: red and cyan).

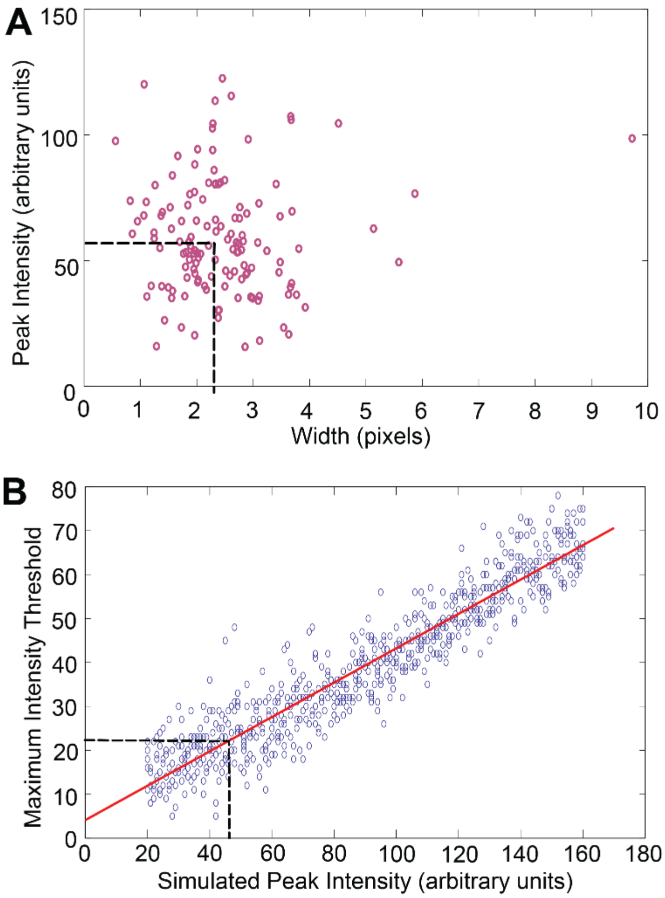


**Figure S5. Determining the number of missed binding events. (A)** Measurement of the width and peak intensity of Cy5-$\sigma^{70}$RNAP fluorescent spots in *experimental* images by Gaussian fitting, yielding median width 2.33 pixels and peak intensity of $I_{exp}$ = 58.6 (dashed lines). B) Maximum value of the intensity threshold setting that allowed detection of a spot of given Gaussian peak intensity above background in a *simulated* image (points) and linear fit (solid line). Dashed lines indicate the threshold setting (21) used in analysis of the experimental data and the corresponding minimum spot intensity detected ($I_{min}$ = 46.2).

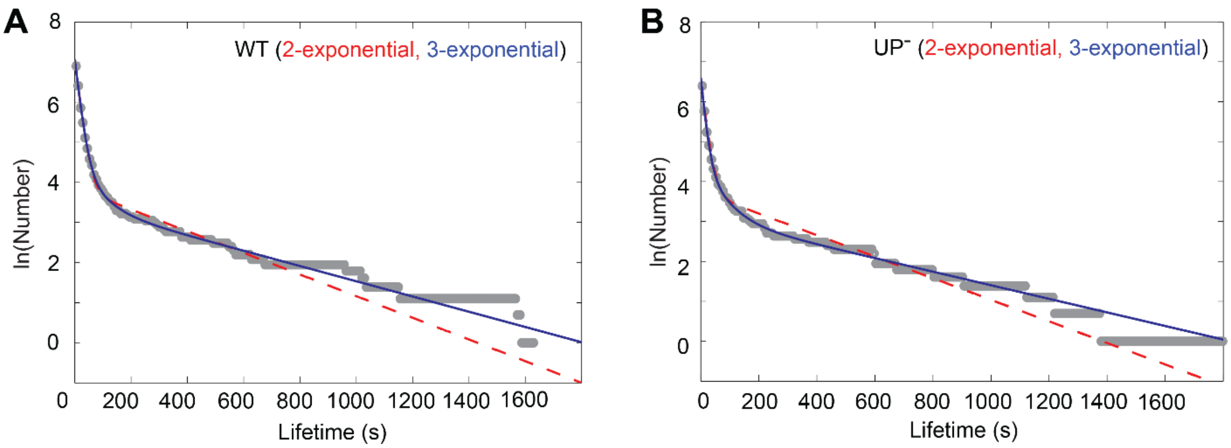


**Figure S6. Measurement and fitting of $\sigma^{70}$RNAP-DNA complex lifetime distributions in the presence of nucleotides.** **(A)** Cumulative distributions (plotted as in Fig. 4) of *rrnB* P1-specific lifetimes (gray) on WT DNA in the presence of 500 µM ATP, CTP, and UTP ($N$ = 990), and the distributions predicted from two- (red) and three-exponential (blue) fits to the data (see Methods and Table S2). **(B)** Same as (A) except using $UP^-$ DNA ($N$ = 598).

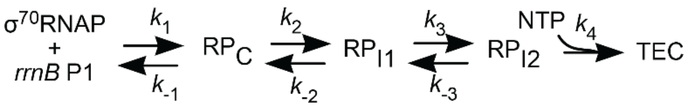


**Figure S7. Hypothetical three-intermediate sequential kinetic pathway of initiation.** This alternative hypothesis was excluded based on the experimental initiation rate and lifetime data (see text).

### Supplemental Tables

#### Table S1. PCR primers and transcript hybridization probes

| Oligonucleotide name | 5' modification | Sequence* |
|---|---|---|
| WT forward† | Alexa488 – $C_{12}$ | 5´-GCGGTCAGAAAATTATTTTAAATTTCC-3´ |
| $UP^{-}$ forward† | Alexa488 – $C_{12}$ | 5´-ATTGACTCCCCGGCGGGGCCCGGGCCTCTTGTCAGGCCGGA-ATAACTCC-3´ |
| $P1^{-}$ forward§ | Alexa488 – $C_{12}$ | 5´-CGGTCAGAAAATTATTTTAAATTTCCTCGGTGACGGCCGGA-ATAACTCCCGCGCCGGCGCCACCACTGACACGGAACAACG-3´ |
| $UP^{-}P1^{-}$ forward§ | Alexa488 – $C_{12}$ | 5´-CATTGACTCCCCGGCGGGGCCCGGGCCTCGGTGACGGCCG-GAATAACTCCCGCGCCGGCGCCACCACTGACACGGAACAACG-3´ |
| *rrnB* reverse§ | Biotin – $C_{12}$ | 5´-CGTGTTCACTCTTGAGACTTGGTATTC-3´ |
| Transcript Probe 1§ | Cy3 – $C_{12}$ | 5´-TGACTGTGCCTTGTTGCCGTTTGTGCG-3´ |
| Transcript Probe 2§ | Cy3 – $C_{12}$ | 5´-GCGGCCCAGTCGCCCCAAGAGGACTCT-3´ |

*Colors show sequence segments containing UP (blue), core promoter elements (red), or their corresponding ablation mutations, and the position of the transcription start site in the WT promoter (green). †From Eurofins. §From IDT DNA.

**Table S2**. Lifetime and amplitude parameters from two- or three-exponential fitting of $\sigma^{70}$RNAP-promoter complex lifetimes.

| DNA | Amp. 1 | $\tau_1$ (s) | Amp. 2 | $\tau_2$ (s) | Amp 3 | $\tau_3$ (s) |
|---|---|---|---|---|---|---|
| **WT −NTP** | 0.92 | 12.9 | 0.08 | 135 | | |
| **± S.E.** | 0.02 | 0.6 | 0.02 | 18 | | |
| **WT +NTP** | 0.96 | 17.3 | 0.04 | 376 | | |
| **± S.E.** | 0.03 | 0.7 | 0.03 | 80 | | |
| **WT −NTP** | 0.00 | 4.7 | 0.914 | 12.9 | 0.086 | 135 |
| **± S.E.** | 0.00 | 0.4 | 0.004 | 0.5 | 0.007 | 20 |
| **WT +NTP** | 0.92 | 16.0 | 0.058 | 69 | 0.026 | 547 |
| **± S.E.** | 0.07 | 0.7 | 0.005 | 31 | 0.0001 | 140 |
| **UP⁻ −NTP** | 0.87 | 8.9 | 0.11 | 64 | 0.014 | 714 |
| **± S.E.** | 0.02 | 0.3 | 0.01 | 10 | 0.003 | 143 |
| **UP⁻ +NTP** | 0.88 | 13.2 | 0.087 | 63 | 0.030 | 588 |
| **± S.E.** | 0.04 | 0.8 | 0.007 | 22 | 0.0001 | 555 |
